# Behavioral and neuronal underpinnings of safety in numbers in fruit flies

**DOI:** 10.1101/629311

**Authors:** Clara H Ferreira, Marta A Moita

## Abstract

Living in a group allows individuals to decrease their defenses enabling other beneficial behaviors such as foraging. The detection of a threat through social cues is widely reported, however the safety cues that guide animals to break away from a defensive behavior and resume alternate activities remain elusive. Here we show that fruit flies displayed a graded decrease in freezing behavior, triggered by an inescapable threat, with increasing group sizes. Furthermore, flies used the cessation of movement of other flies as a cue of threat and its resumption as a cue of safety. Finally, we found that lobula columnar neurons, LC11, mediate the propensity for freezing flies to resume moving in response to the movement of others. By identifying visual motion cues, and the neurons involved in their processing, as the basis of a social safety cue this study brings new insights into the neuronal basis of safety in numbers.

## Introduction

Predation is thought to be a key factor driving group formation and social behavior (reviewed in (Alexander, 1974)). It has long been established that being in a group can constitute an anti-predatory strategy (Foster and Treherne, 1981; Hamilton, 1971), as it affords the use of social cues to detect predators (Murray et al., 2017; Pereira et al., 2012), enables coordinated defensive responses (Ono and Sasaki, 1987) or simply dilutes the probability of each individual to be predated (Foster and Treherne, 1981). A major consequence of the safety in numbers effect, reported in taxa throughout the animal kingdom, is that animals tend to decrease their individual vigilance (Underwood, 1982), stress levels (Queiroz and Magurran, 2005) or defensive behaviors (Faustino et al., 2017) when in a social setting. Despite its wide prevalence the mechanisms that lead to a decrease in defensive behaviors are largely unknown. Hence, in order to gain mechanistic insight into how increasing group size impacts defense behaviors, we decided to use *Drosophila melanogaster* since it allows the use of groups of varying size, the large number of replicates required for detailed behavioral analysis and genetic access to specific neuronal subtypes. Importantly, fruit flies display social behaviors in different contexts (Battesti et al., 2012; Combes et al., 2012; Danchin et al., 2018; Kacsoh et al., 2015; Ramdya et al., 2015; Sarin and Dukas, 2009), namely social regulation of anti-predation strategies, such as the socially transmitted suppression of egg laying in the presence of predatory wasps (Sarin and Dukas, 2009) or the reduction in erratic turns during evasive flights when in a group, compared to when alone, in the presence of dragonflies (Combes et al., 2012).

## Results

### Flies in groups display lower sustained freezing responses

To simulate a predator’s attack, we used a looming stimulus (Fig. 1A), an expanding dark disc, that mimics an object on collision course and elicits defense responses in visual animals, including humans (reviewed in (Fotowat and Gabbiani, 2011; Herberholz and Marquart, 2012; Peek and Card, 2016)). Individually tested fruit flies respond to looming stimuli with escapes in the form of jumps (Card and Dickinson, 2008; Reyn et al., 2014), in flight evasive maneuvers (Muijres et al., 2014) or running as well as with freezing (Gibson et al., 2015; Zacarias et al., 2018) when in an enclosed environment. In our setup, the presentation of 20 looming stimuli (Fig. 1A) elicited reliable freezing responses for flies tested individually and in groups of up to 10 individuals (Fig. 1B-E, Supplemental Fig. 1 shows that running and jumps are less prominent in these arenas). The fraction of flies freezing increased as the stimulation period progressed for flies tested individually and in groups of up to 5 flies; in groups of 6 to 10 individuals, the fraction of flies freezing only transiently increased with each looming stimulus (Fig. 1B). The fraction of flies freezing was maximal for individuals and minimal for groups of 6 to 10, while groups of 2 to 5 flies showed intermediate responses (Fig. 1B). At the level of each individual fly’s behavior, flies tested alone spent more time freezing, 76.67%, IQR 39.75-90.42%, during the stimulation period than flies in any of the groups tested (Fig. 1C; statistical comparisons in Supplemental Table 1). Flies in groups of 2 to 5 spent similar amounts of time freezing (for groups of 2: 31.67%, IQR 9.46-64.38% and for groups of 5: 43.08%, IQR 11.79-76.50%), while flies in groups of 6 to 10 displayed the lowest levels of freezing (for groups of 6: 8.08%, IQR 3.04-17.46% and for groups of 10: 3.33%, IQR 2-7.67%) (Fig. 1C; statistical comparisons in Supplemental Table 1). The decrease in time spent freezing for flies tested in groups of 2 to 5, compared to individuals, was not due to a decrease in the probability of entering freezing after a looming stimulus (Fig. 1D; statistical comparisons in Supplemental Table 2), but rather to an increase in the probability of stopping freezing, i.e. resuming movement, before the following stimulus presentation (individually tested flies: P(F_exit_)=0.08, IQR 0-0.21, groups of 2: P(F_exit_)=0.31 IQR 0.11-0.78, groups of 5: P(F_exit_)=0.54 IQR 0.31-0.90) (Fig. 1E; statistical comparisons in Table S3). Flies in groups of 6 to 10, were not only more likely to stop freezing (groups of 6: P(F_exit_)=0.93, IQR 0.80-1, groups of 10: P(F_exit_)= 1, IQR 0.83-1) (Fig. 1E; statistical comparisons in Table S3), but also less likely to enter freezing (groups of 6: P(F_entry_)=0.35, IQR 0.20-0.46, groups of 10: P(F_entry_)= 0.21, IQR 0.10-0.36) (Fig. 1C; statistical comparisons in Supplemental Table 2) compared to the other conditions. The decrease in persistent freezing with the increase in group size suggests that there is a signal conveyed by the other flies that increases in intensity with the increase in the number of flies tested together.

**Fig. 1.**
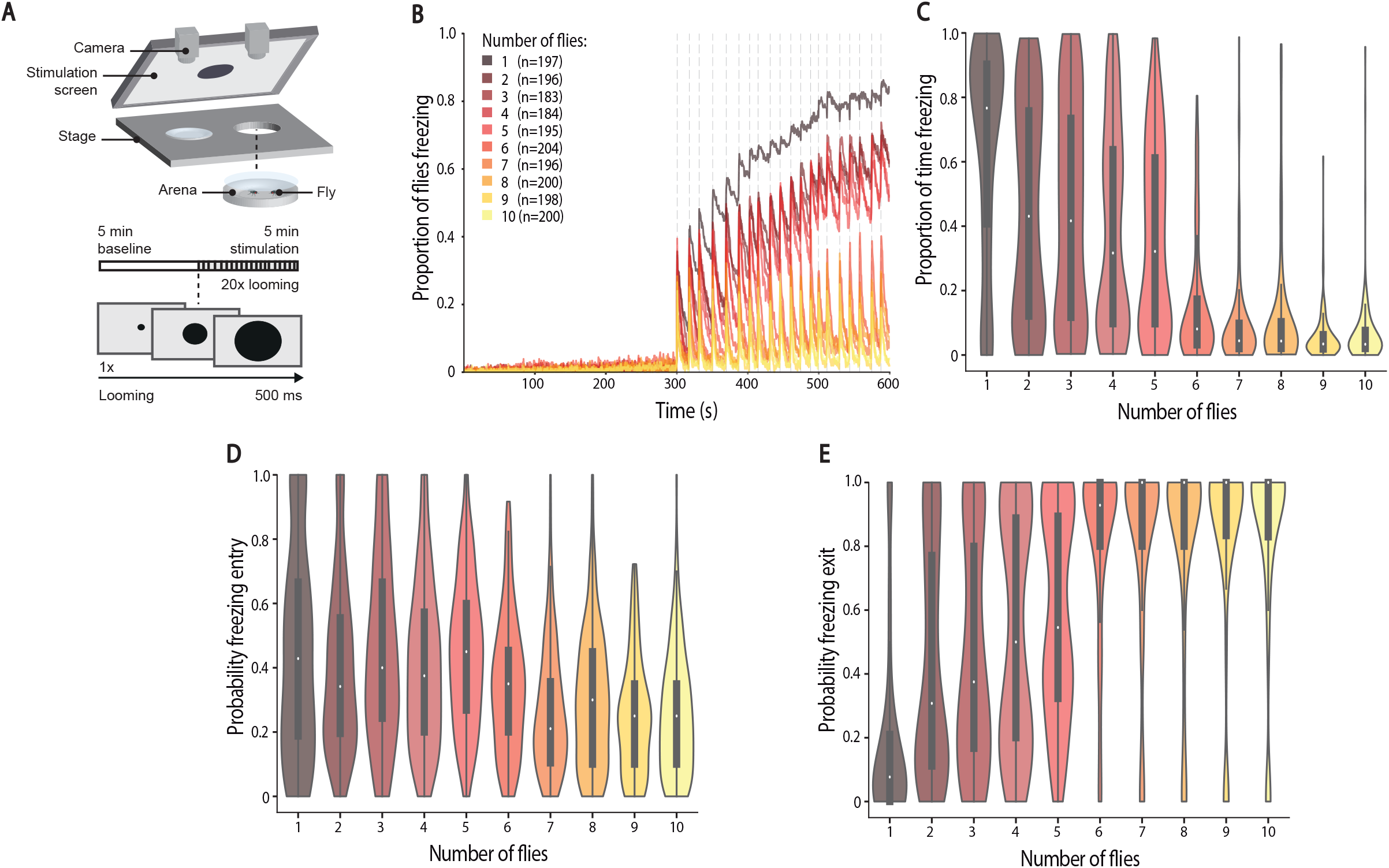
Analysis of the group effect on freezing responses. A) Experimental setup and protocol. We tested individuals and groups of up to 10 flies in backlit arenas imaged from above. After a 5-minute baseline flies were exposed to twenty 500 ms looming presentations, every 10-20 s, indicated by vertical dashed lines. B) Proportion of flies freezing throughout the experiment. C–E) Violin plots representing the probability density of individual fly data bound to the range of possible values, with boxplots. C) Proportion of time spent freezing throughout the experiment. Statistical comparisons between conditions presented in Table S1. D) Probability of freezing entry in the 500 ms bin following looming presentation. Statistical comparisons between conditions presented in Table S2. E) Probability of freezing exit in the 500 ms bin before the following looming stimulus. Statistical comparisons between conditions presented in Table S3.

### Absence of movement promotes freezing

We next examined whether flies respond to each other. We started by exploring the effect on freezing onset, as freezing has been shown to constitute an alarm cue in rodents, such that one rat freezing can lead another to freeze (Pereira et al., 2012). We decided to focus on groups of 5 flies, which showed intermediate freezing levels (Fig. 1). The onset of freezing both for individually tested flies and in groups of 5 occurred during and shortly after a looming stimulus (Fig. 2A). This window, of ~1s, in principle allows for social modulation of freezing onset. Indeed, the probability of freezing onset at time *t* gradually increased with increasing numbers of flies freezing at time *t-1* (see methods), indicating that flies increase their propensity to freeze the more flies around them were freezing. This synchronization in freezing could result from flies being influenced by the other flies or simply time locking of freezing to the looming stimulus. To disambiguate between these possibilities we shuffled flies across groups, such that the virtual groups thus formed were composed of flies that where not together when exposed to looming. If the looming stimulus was the sole source of synchrony for freezing onset, then we should see a similar increase in probability of freezing by the focal fly with increasing number of ‘surrounding’ flies freezing in the shuffled group. We found a weaker modulation of freezing onset by the number of flies freezing in randomly shuffled groups compared to that of the real groups of 5 flies (Fig. 2B; G-test, g=190.96, p<0.0001, df=4). We corroborated this result by testing single flies surrounded by 4 fly-sized magnets whose speed and direction of circular movements we could control (Fig. 2C-F). During baseline, the magnets moved at the average walking speed of flies in our arenas, 12 mm/s, with short pauses as the direction of movement changed. Stopping the magnets upon the first looming stimulus and throughout the entire stimulation period led to increased time freezing (Fig. 2D) and increased probability of freezing entry upon looming (Fig. 2E), compared to all controls – individuals alone, magnets not moving throughout the entirety of the experiment and the exact same protocol (magnets moving during baseline then freezing) but in the absence of looming stimuli. The transition from motion to freezing is thus important, but not sufficient to drive freezing, since flies surrounded by magnets that do not move for the entire experiment froze to individually-tested levels, but flies exposed to magnets that move and then freeze in the absence of looming stimuli did not freeze. Together these results suggest that flies use freezing by others as an alarm cue, which increases their propensity to freeze to an external threat, the looming stimulus.

**Fig. 2.**
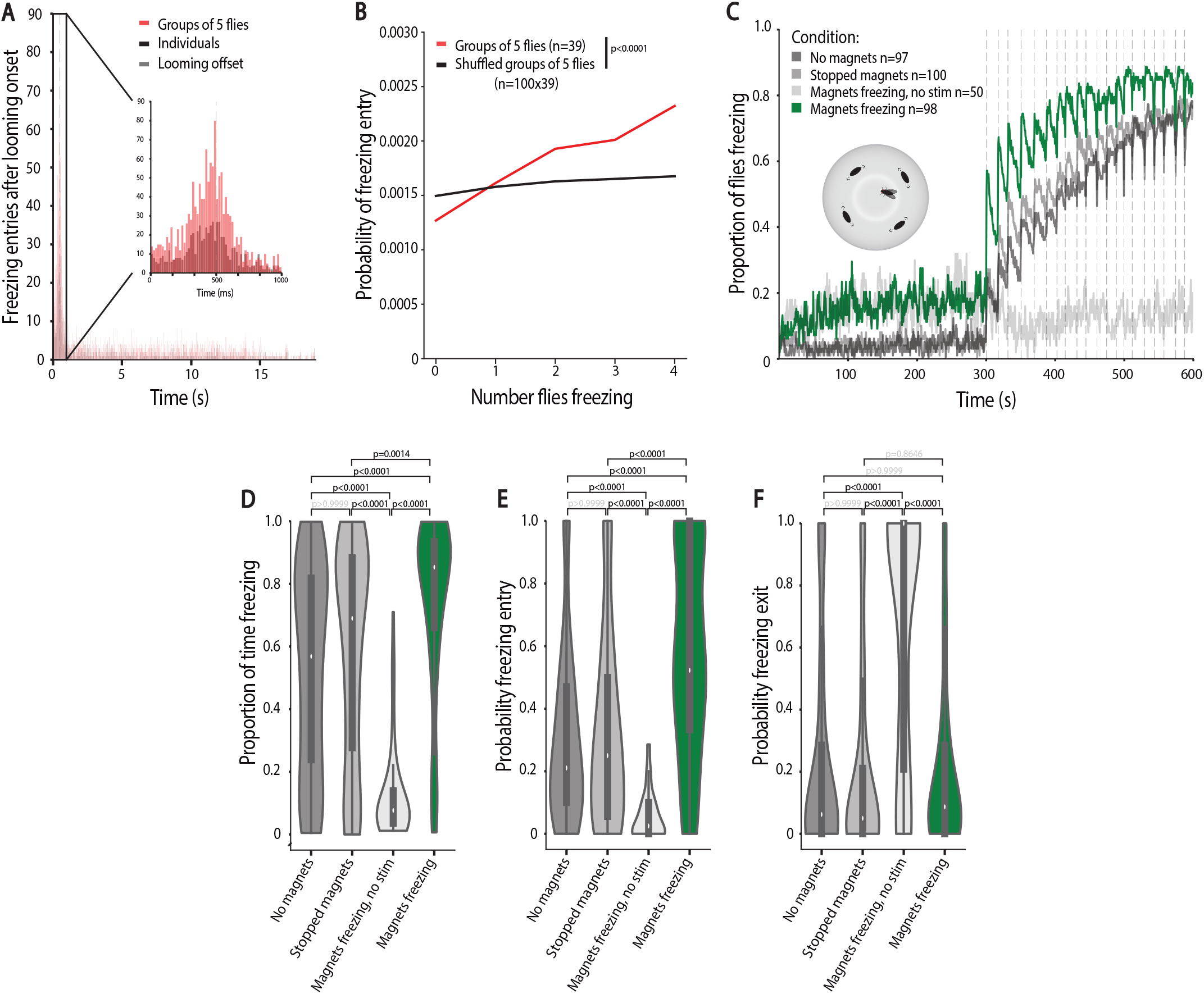
The group effect on individual freezing entry. A) Distribution of freezing entries after looming onset for flies tested individually and in groups of 5. B) Probability of freezing entry at time t as a function of the number of other flies freezing at t-1 (see methods). C-F) Simulating groups of 5 using movable magnets. C) Proportion of flies freezing throughout the experiment. D–F) Violin plots representing the probability density of individual fly data bound to the range of possible values, with boxplots. D) Proportion of time spent freezing throughout the experiment. E) Probability of freezing entry in the 500 ms bin following looming presentation. F) Probability of freezing exit in the 500 ms bin before the following looming stimulus. P-values result from Kruskal-Wallis statistical analysis followed by Dunn’s multiple comparisons test.

### Movement of neighbors leads to freezing exit and movement resumption

As the strongest effect observed across all group sizes was on freezing exit, i.e. the resumption of movement, we asked whether the propensity to exit freezing was also dependent on the number of surrounding flies that were freezing. To this end, we performed a similar analysis as for freezing onset and found that the higher the number of flies freezing the lower the probability of the focal fly to exit from freezing. This effect was also decreased in shuffled groups (Fig. 3A; G-test, g=170.81, p<0.0001, df=4). We then examined the contribution of mechanosensory signals in the decrease in freezing and found that collisions between flies play a minor role in the observed effect (Supplemental Fig. 2; statistical comparisons in Supplemental Tables 4-6), contrary to what happens with socially-mediated odor avoidance (Ramdya et al., 2015). Next, we explored our intuition that motion cues from the other flies were the main players affecting exit from looming-triggered freezing. We formalized the motion signal (Fig. 3B), perceived by a focal fly, as the summed motion cues produced by the other four surrounding flies (we multiplied the speed of each fly by the angle on the retina, a function of the size of the fly and its distance, to the focal fly, Fig. 3B). We then analyzed separately the summed motion cue perceived by focal flies during freezing bouts that terminated before the following looming stimulus (freezing with exit) and continuous freezing bouts (with no breaks in between looming stimuli) (representative examples in Fig. 3B). Freezing bouts with exit had higher motion cue values (Fig. 3C) compared to continuous bouts (p<0.0001, Freezing without exit=0.64 IQR: 0.00-2.11, Freezing with exit=2.79 IQR: 1.28-5.08).

**Fig. 3.**
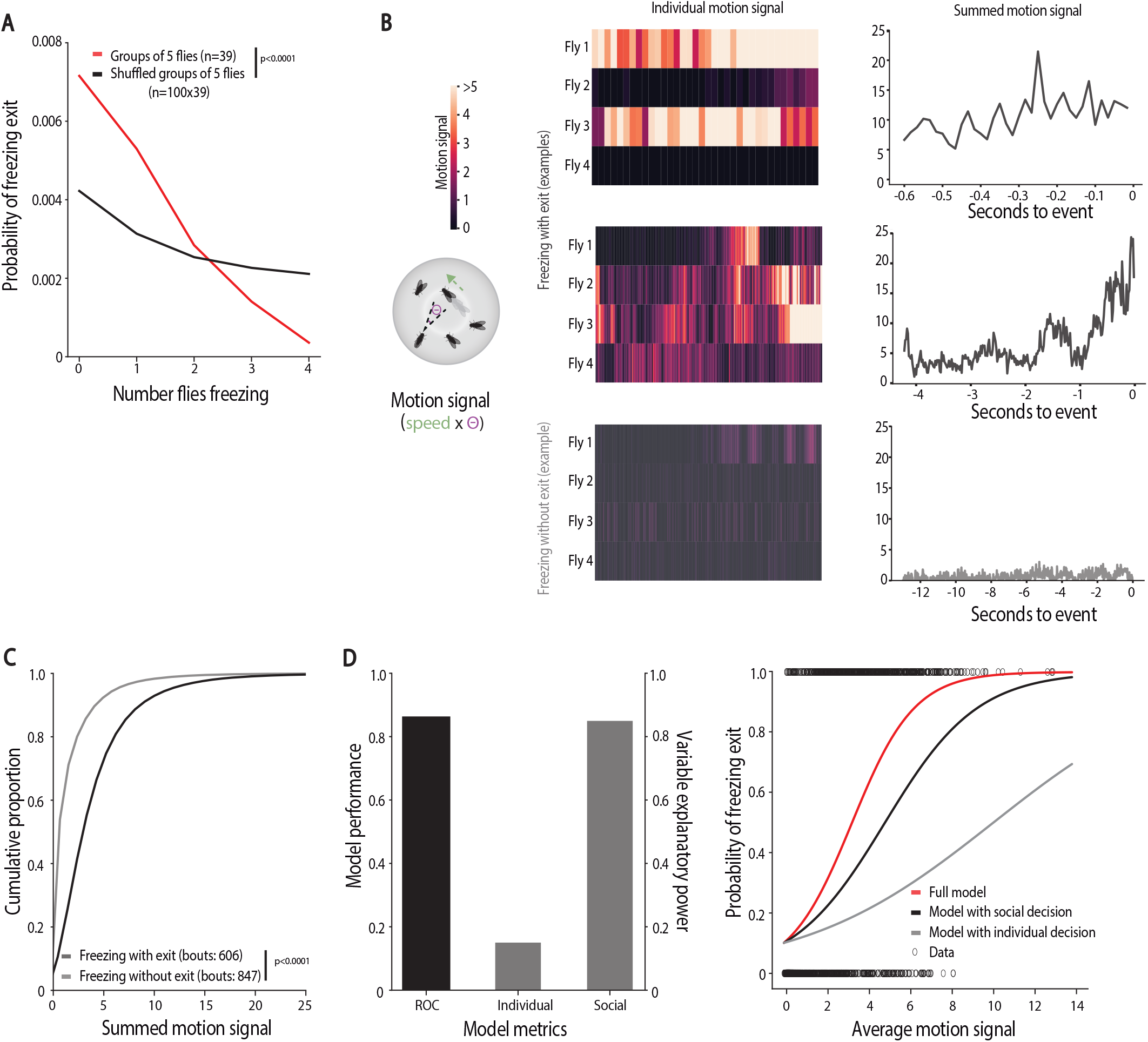
The group effect on individual freezing exit. A) Probability of freezing at time t as a function of the number of flies freezing at time t-1 (see methods). B) The motion signal is formalized as the other fly’s speed multiplied by the angle it produces on the retina of the focal fly (schematic). Representative examples of the motion signal starting in the 500 ms bin after looming offset by a focal fly until freezing exit or the end of the inter-looming interval (without freezing exit): heatmaps show the motion signals of the 4 surrounding flies and the line graphs show the summed motion signal. C) Cumulative distributions of the summed motion signals. D) Logistic regression model of the decision to stop or continue freezing as a function of individual and social decision processes. Left panel: model performance and explanatory power of the individual and social processes; right panel: binary freezing data and logistic predictive probabilities using individual, social variables and coefficients alone or combined as a function of the average motion signal.

We hypothesized that once flies start freezing, upon a looming stimulus, two processes determine whether a fly will exit freezing, resuming activity, or remain freezing: 1) an individual decision process, whereby flies make this binary decision irrespective of what the other flies are doing, possibly reflecting the number of looming stimuli the flies were exposed to and how much time has elapsed since the onset of freezing; 2) a social decision process whereby flies integrate the motion cues generated by their neighbors relying on this information to decide whether to stop freezing. To test this possibility, we modeled the decision to stay freezing or resume activity as a binary decision that follows a logistic function taking into account two parameters, the individual probability of exiting freezing before the next looming stimulus, and the motion cues of others (see methods). With this simple model we can predict whether a fly will stay freezing during the entire inter-looming interval or whether it resumes activity in between looming stimuli, (ROC = 0.86, Fig. 3D). In addition, we found that the social cues explained a large fraction of the variance while individual behavior explains a small fraction (average variance explained by β-coefficient of social cues, β_s_ = 0.85, variance explained by β-coefficient for individual behavior β_i_ = 0.15, Fig. 3D).

To further test whether motion cues from others constitute a safety signal, we manipulated the motion cues perceived by the focal fly, while maintaining the number of flies in the group constant. An increase in the social motion cues, should enhance the group effect, and hence decrease the freezing responses of a focal fly. We compared groups of 5 wild-type flies with groups of 1 wild-type and 4 blind flies (Fig. 4A). Blind flies don’t perceive the looming stimulus and walk for the duration of the experiment; when a focal fly freezes surrounded by 4 blind flies it is thus exposed to a higher motion signal during the stimulation period than a focal fly in a group of 5 wild-type flies (Fig. 4A). When surrounded by blind flies, the fraction of focal flies freezing throughout the stimulation period was lower than the fraction of flies freezing in a group of wild-type flies (Fig. 4B). Further, the increase in motion cues in groups with blind flies decreased the amount of time a fly froze compared to that of groups of wild-type flies (6.17% IQR 2.17-15.25% versus 19.58% IQR 8.20-57.12; p<0.0001) (Fig. 4C). This reduction in freezing resulted mostly from a decreased probability of freezing entry (wild-type groups: P(F_entry_)=2.57 IQR 0.15-0.39, groups with blind flies: P(F_entry_)=0.49 IQR 0.25-0.61, p<0.0001) (Fig. 4D) and slightly increased probability of exiting freezing (wild-type groups: P(F_exit_)= 0.83 IQR 0.39-1, groups with blind flies: P(F_exit_)= 0.89 IQR 0.71-1) (Fig. 4E). Hence, a focal fly surrounded by 4 blind flies behaves similarly to flies in groups of more than 6 individuals. Importantly, the decrease in persistent freezing was not due to an increased role of collisions on freezing breaks (Supplemental Fig. 3). We further tested whether any type of visual signal could alter individual freezing in the same manner as the motion cues generated by flies in the group, by presenting a visual stimulus with randomly appearing black dots with the same change of luminance as the looming stimulus but without motion. (as in *19*) 4.5 seconds after each looming presentation. This stimulus, which could work as a distractor, did not alter the proportion of time freezing nor the probability of freezing entry or exit (Supplemental Fig. 3).

**Fig. 4.**
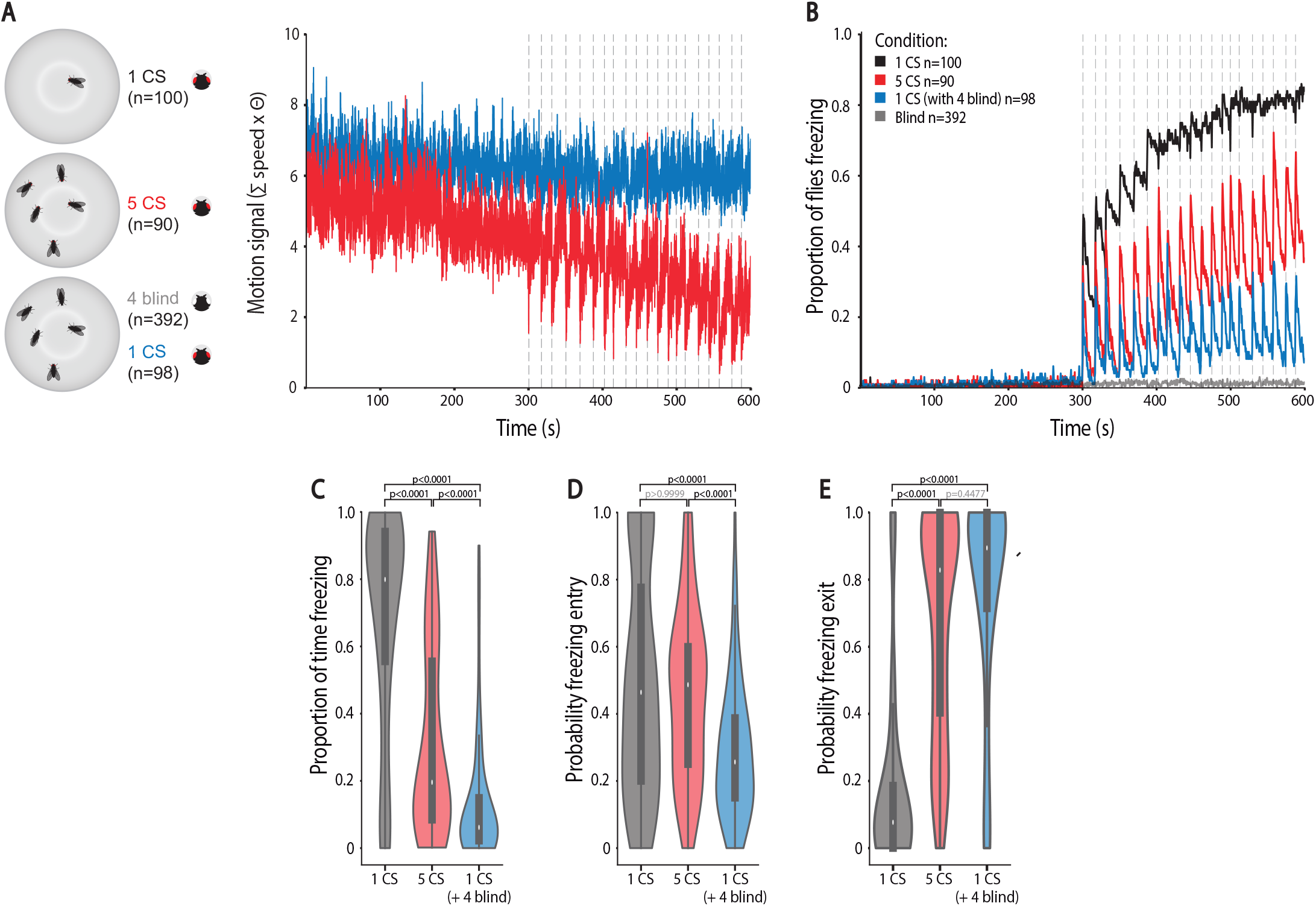
Manipulating the motion signal in groups of 5 flies. A) The summed motion signal of surrounding flies for groups of 5 wild-type flies and groups with one wild-type and 4 blind flies. B) Proportion of flies freezing throughout the experiment. C–E) Violin plots representing the probability density of individual fly data bound to the range of possible values, with boxplots. C) Proportion of time spent freezing throughout the experiment. D) Probability of freezing entry in the 500 ms bin following looming presentation. E) Probability of freezing exit in the 500 ms bin before the following looming stimulus. P-values result from Kruskal-Wallis statistical analysis followed by Dunn’s multiple comparisons test.

Together, these results show that flies use motion cues generated by their neighbors to decide whether to stay or exit freezing, raising the possibility that motion cues produced by others could constitute a safety signal leading flies to resume activity.

### Lobula columnar neurons 11 mediate group effect on freezing responses

Having identified motion cues of others as the leading source of the group effect on freezing, we decided to test the role of visual projection neurons responsive to the movement of small objects, lobula columnar 11 (LC11) (Keleş and Frye, 2017; Wu et al., 2016) in our paradigm. The behavioral relevance of these neurons was as yet unidentified. We used two fly lines, an *LC11-GAL4* (Keleş and Frye, 2017) and an *LC11-splitGAL4* (Wu et al., 2016), to drive the expression of Kir 2.1 (Baines et al., 2001), a potassium channel that hyperpolarizes neurons. Anatomically, both fly lines encompass LC11 neurons (Fig. 5A and B) but *LC11-splitGAL4* is more specific as it does not contain the neurons in the subesophageal zone that descend to the thoracico-abdominal ganglion present in the *LC11-GAL4*. Constitutively expressing Kir 2.1 in LC11 neurons did not alter looming-triggered freezing of flies tested individually, when compared to parental controls (Supplemental Fig. 4). Conversely, for LC11-silenced flies tested in groups of five, the fraction of flies freezing increased throughout the experiment (Fig. 5A and B). Moreover, experimental flies in groups of 5 froze longer (~6 fold increase for *LC11-GAL4 >Kir2.1*, and ~2 and 7 fold increase for *LC11-splitGAL4 >Kir2.1* compared to parental controls) (Fig. 5C), which was not due to an increase in the probability of freezing entry (Fig. 5D), but rather to a decrease in the probability of freezing exit (Fig. 5E) (*LC11-GAL4 >Kir2.1* 0.12 IQR 0.05-0.50, *LC11-GAL4/+* 0.786 IQR 0.44-1 and *Kir2.1/+* 0.83 IQR 0.5-1, p<0.; *LC11-splitGAL4 >Kir2.1* 0.24 IQR 0.06-0.67, *LC11-splitGAL4/+* 1 IQR 0.67-1 and *Kir2.1/+* 0.50 IQR 0.26-0.86, p=0.0001). LC11 neurons have been shown to be maximally responsive to moving objects with an angular size of 2.2° and display half amplitude responses to a 4.4° object (Keleş and Frye, 2017) corresponding, in our arenas, to stronger LC11 responses to moving flies further away (a fly at the maximal possible distance, 6.5 cm has an angular size of ~2.6° while flies with a 4.4° angular size would be at a distance of ~3.4 cm from a focal fly). Interestingly, silencing these neurons did not affect the use of freezing as an alarm cue, since these flies showed an increased probability of freezing the more surrounding flies were freezing (Supplemental Fig. 4). Finally, expressing Kir2.1 in another LC neuron class, LC20 (Wu et al., 2016), which are not known to respond to small moving objects, does not alter group behavior (Supplemental Fig. 5). In summary, silencing LC11 neurons renders flies less sensitive to the motion of others, specifically decreasing its use as a safety cue that downregulates freezing.

**Fig. 5.**
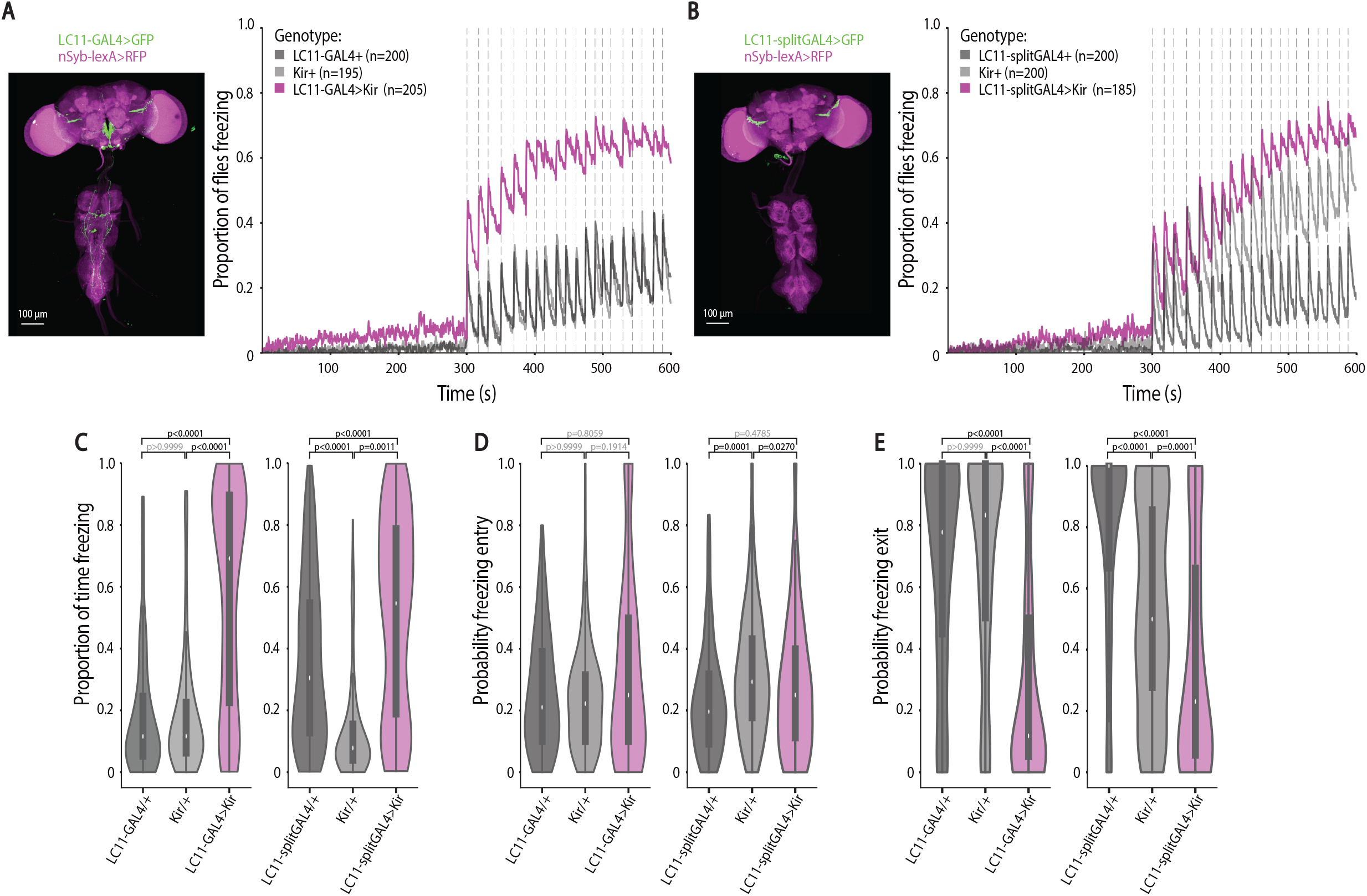
Manipulating lobula columnar neurons 11 (LC11). A–B) Anatomy (scale bars, 100 μm) and proportion of flies freezing throughout the experiment in groups of 5, for A) *LC11-GAL4>Kir2.1* and B) *LC11-splitGAL4>Kir2.1* depicted in purple and parental controls (grey). C–E) Violin plots representing the probability density distribution of individual fly data bound to the range of possible values, with boxplots. C) Proportion of time spent freezing throughout the experiment. D) Probability of freezing entry in the 500 ms bin following looming presentation. E) Probability of freezing exit in the 500 ms bin before the following looming stimulus. P-values result from Kruskal-Wallis statistical analysis followed by Dunn’s multiple comparisons test.

## Discussion

In this study we show that flies in groups display a reduction in freezing responses that scales with group size. Detailed behavioral analysis and quantitative modeling together with behavioral and genetic manipulations, allowed us to identify freezing as a sign of danger and activity as a safety cue. These findings are consistent with the hypothesis that safety in numbers may partially be explained by the use of information provided by the behavior of others. Moreover, we show that visual projection LC11 neurons are involved in processing motion cues of others to down regulate freezing.

Because living in a group allows individuals to decrease their defenses, it also enables other globally beneficial behaviors such as foraging. These selective forces on the evolution of social behavior have been demonstrated in a wide range of animals ranging from invertebrates to mammals (Alexander, 1974; Hamilton, 1971). One of the main benefits of being in a group is the facilitated detection of behaviourally significant cues in the environment, as information about their presence can quickly spread across a large group of individuals (Handegard et al., 2012). In the context of threat detection, most research has focused on actively emitted signals, such as alarm calls and foot stamping (reviewed in (Pereira and Moita, 2016; Rose et al., 2006)). However, cues generated by movement patterns produced by defensive responses of surrounding prey can play a crucial role in predator detection. For example, crested pigeons use distinct wing whistles produced by conspecific escape flights (Murray et al., 2017) and rats use silence resulting from freezing, as alarm cues (Pereira et al., 2012). Recently, it has also been suggested that seismic waves produced by fast running in elephants promote vigilance in conspecifics(Mortimer et al., 2018). This form of social detection of threat may be advantageous as it does not require the active production of a signal that may render the emitter more conspicuous and thus vulnerable. Although few studies demonstrated this phenomenon, it is described in distant vertebrate species. In this study, we extend to invertebrates the notion of defensive behaviors, in this case freezing, as alarm cues. In addition, freezing may constitute a public cue that can be used by any surrounding animal regardless of species. Indeed, we show that freezing by ‘dummy’ flies enhances freezing in response to looming stimuli.

Importantly, we also identify a social cue of safety. In our paradigm flies responded to the threatening looming stimulus with freezing. At some point after the stimulus, flies can exit freezing resuming movement, until a new looming stimulus is presented, triggering freezing again. The more stimuli the flies were exposed to the less likely they are to exit freezing before the next looming. This pattern suggests that the resumption of activity reflects the level of safety, such that when in groups the movement of others can constitute a cue of safety leading to further activity. Using a logistic regression model and manipulating the levels of movement by neighboring flies we demonstrated that motion cues of others strongly determine the propensity of flies to resume activity. In a prior study (Pereira et al., 2012) we show that when we present an auditory cue of movement to rats that are freezing in response to the display of freezing by another rat, they resume activity. Although in line with the present findings, we did not explicitly test whether this motion cue constituted a safety cue, as we have done here.

While the known examples of the use of auditory motion cues to infer the presence or absence of a threat in vertebrate species, here we show that flies use visual motion cues. This may relate to the fact that insects can only use short range auditory signals, whereas visual cues can be detected at larger distances. Silencing LC11 neurons, which process visual information, responding to motion of small visual objects, disrupted the use of motion cues from neighboring flies as a safety cue. Interestingly, the role of these cells was quite specific, silencing them did not affect the use of freezing as an alarm cue. Furthermore, other LC neurons have been implicated in processing visual stimuli in social contexts, namely *fru+* LC10a important for the ability of males to follow the female during courtship (Ribeiro et al., 2018). In mammals, there are roughly 20-30 distinct retinal ganglia cell (RGC) types, and only recently has work begun to detail the link between the features that distinct RGCs detect, and the specific behaviors their activity promotes (reviewed in (Sanes and Masland, 2015). LC cells in the fly seem to have a similar organization (reviewed in (Joly et al., 2016; Sanes and Zipursky, 2010)), wherein LC neurons are tuned for distinct visual features, and activating specific LC cells leads to distinct approach or defensive responses. It will be interesting to study to what extent there is specificity or overlap in visual projection neurons for behaviors triggered by the motion of others. The parallels between visual systems of flies and humans, despite the lack of any common ancestor with an image forming visual system, suggest that shared mechanisms underlying visuomotor transformations represent general solutions to common problems that all organisms face individually or as a group.

Motion plays a crucial role in predator-prey interactions. Predator and prey both use motion cues to detect each other using these to make decisions about when and how to strike or whether and how to escape (Carr and Christensen-Dalsgaard, 2015; Catania et al., 2008; Friedel et al., 2008; Zhao et al., 2019). Furthermore, prey animals also use motion cues from other prey as an indirect cue of a predator’s presence (Handegard et al., 2012; Hingee and Magrath, 2009; Pereira et al., 2012). We believe that the current study opens a new path to study how animals in groups integrate motion cues generated by predators, their own movement, and that of others to select the appropriate defensive responses.

## Acknowledgments

We would like to thank: the Scientific Software Platform at the Champalimaud Centre for Unknown for developing the Fly motion quantifier; the Scientific Hardware platform for developing the magnet setup; Wolf Huetteroth (University of Leipzig) for help with imaging fly lines; Ricardo Vieira for help streamlining the video analysis pipeline; Gil Costa for the illustrations in Figure 1A and Figure 3B and 4A; the Moita lab, particularly Anna Hobbiss and Ricardo Neto, as well as Eugenia Chiappe and Gonzalo de Polavieja for fruitful discussions and comments on the manuscript; Alfonso Renart and João Afonso for help with the logistic regression model. Funding: This work was supported by Fundação Champalimaud, ERCStG337747-CoCO and ERCCoG819630-A-Fro.

## Author contributions

C.H.F. performed all experiments and analyzed the data. C.H.F. and M.A.M. designed the experiments, discussed results and wrote the manuscript. Authors declare no competing interests.

## Data availability

Data and materials available upon request.

## Material and methods

### Fly lines and husbandry

Flies were kept at 25 °C and 70% humidity in a 12 h:12 h dark:light cycle. Experimental animals were mated females, tested only once when 4–6 days old.

Wild-type flies used were Canton-S. LC11-splitGAL4 line *w[1118]; P{y[+t7.7] w[+mC]=R22H02-p65.AD}attP40; P{y[+t7.7] w[+mC]=R20G06-GAL4.DBD}attP2*, LC11-GAL4 w[1118]; P{y[+t7.7] w[+mC]=GMR22H02-GAL4}attP2 and LC20-splitGAL4 *w[1118]; P{y[+t7.7] w[+mC]=R35B06-GAL4.DBD}attP2 PBac{y[+mDint2] w[+mC]=R17A04-p65.AD}VK00027* and w[*] norpA[36] were obtained from the Bloomington stock center. *10XUAS-IVS-eGFPKir2.1 (attP2)* flies were obtained from the Card laboratory at Janelia farm. UAS-CD8::GFP; lexAop-rCD2::RFP (Lee et al., 1999) recombined with nSyb-lexA.DBD::QF.AD (obtained from the Bloomington stock center) were obtained from Wolf Huetteroth (University Leipzig).

### Behavioral apparatus and visual stimulation

We imaged unrestrained flies in 5 mm thick, 11° slanted polyacetal arenas with 68 mm diameter (central flat portion diameter 32 mm). Visual stimulation (twenty 500 ms looming stimuli, a black circle in a white background, with a virtual object length of 10 mm and speed 25 cm/s (l / v value of 40 ms) as in (Zacarias et al., 2018)) was presented on an Asus monitor running at 144 Hz, tilted 45° over the stage (Figure 1A). For the experiments with random dots, 4.5 s after the looming presentation we presented a visual stimulus consisting of appearing black dots at random locations on the screen to reach the same change in luminance as the looming stimulus (Zacarias et al., 2018).

The stage contained two arenas, backlit by a custom-built infrared (850 nm) LED array. Videos were obtained using two USB3 cameras (PointGrey Flea3) with an 850 nm long pass filter, one for each arena.

For the experiments with the magnets (Figure 2), we used an electromechanical device developed by the Scientific Hardware Platform at the Champalimaud Centre for the Unknown. It consists of an adapted setup in which a rotating transparent disc with 5 incorporated neodymium magnets moves under the arena. A circular movement is induced by an electric DC gearhead motor transmitted via a belt to the disc. This allows the movement of magnetic material placed on the arena to move around in synchronized motion. The motor is controlled by a custom-made electronic device, connected to the computer and using a dedicated Champalimaud Hardware Platform software. For the experiments of freezing magnets during stimulation, with or without stimulus, the magnets rotated at 12 mm/s with a change in direction every 50 s during the baseline; as soon as the stimulation period started, in synchrony with the first looming stimulus, the magnets ceased movement, until the end of the experiment.

### Video acquisition and analysis

Videos were acquired using Bonsai (Lopes et al., 2015) at 60 Hz and 1280 width × 960 height resolution. We used IdTracker (Pérez-Escudero et al., 2014) to obtain the position throughout the video of each individual fly. The video and the IdTracker trajectories file were then fed to the ‘Fly motion quantifier’, developed by the Scientific Software Platform at the Champalimaud Centre for the Unknown (https://bitbucket.org) in order to obtain the final csv file containing not only position and speed for each fly, but also pixel change in a region of interest (ROI) around each fly defined by a circle with a 30 pixel radius around the center of mass of the fly.

### Data analysis

Data were analyzed using custom scripts in spyder (python 3.5). Statistical testing was done in GraphPad Prism 7.03, and non-parametric, Kruskal-Wallis followed by Dunn’s multiple comparison test or two-tailed Mann-Whitney tests were chosen, as data were not normally distributed (Shapiro-Wilk test). Probabilities were compared using the χ^2^ contingency test in python (G-test).

Freezing was classified as 500 ms periods with a median pixel change over that time period < 30 pixels within the ROI. The proportion of time spent freezing was quantified as the proportion of 500 ms bins during which the fly was freezing.

We calculated the proportion of freezing entries upon looming and exits between looming stimuli (Figure 1) using the following definitions: 1) freezing entries corresponded to events where the fly was not freezing before the looming stimulus (a 1 s time window was used) and was freezing in the first 500 ms bin after the looming stimulus; 2) freezing exits were only considered if sustained, that is when the fly froze upon looming but exited from freezing and was still moving by the time the next looming occurred, i.e. the first 500 ms bin after looming the fly was freezing and in the last 500 ms bin before the next looming the fly was not freezing.

To determine the time of freezing onset or offset (Figure 2A, B and Figure 3A), we used a rolling window of pixel change (500 ms bins sliding frame by frame) and the same criterion for a freezing bin as above). Time stamps of freezing onset and offset were used to calculate the probability of entering and exiting freezing as a function of the number of flies freezing. For freezing entries after looming as well as probabilities of entering and exiting freezing, we considered only instances in which the preceding 500 ms bin was either fully non-freezing or freezing. To determine the numbers of others freezing at freezing entry or exit we used a 10 frame bin preceding the freezing onset or offset timestamp.

Distances between the center of mass of each fly were calculated using the formula 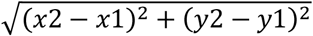, and we considered a collision had taken place when the flies reached a distance of 25 pixels. Motion signal was determined as Σ *speed* × *angle on the retina* (*θ*) where 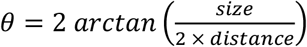.

To analyze the motion signal for freezing bouts with and without exit (Figure 3B, C), we defined freezing bouts with exit as bouts where flies were freezing in the 500 ms following the looming stimulus offset and resumed moving before the next looming stimulus (up until the last 500 ms before the looming stimulus onset) and freezing bouts without exit as those where freezing persisted until the next looming.

To model the decision to stay freezing or resume movement we used the scikit-learn logistic regression model. Briefly, we analyzed freezing behavior in between looming stimuli, categorizing freezing bouts into two types: freezing bouts that ended with an exit before the next looming (to which we assigned a value of 1), and continuous freezing bouts, without an exit until the next looming (value of 0). We used freezing bout type as the dependent variable. The independent variables were the probability of an individual fly exiting from freezing within the same inter-looming interval (calculated from the data of flies tested individually) (V_i_); and the sum of the motion signal generated by neighboring flies, divided by the bout length (V_s_). We performed a K-fold cross-validation with 4 splits. To determine the explanatory power of each predictor, we determined the associated fraction of variance using the following formula (shown for variable V_i_): 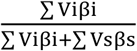.

### Imaging

LC11-GAL4>UAS-CD8::GFP; nSyb-lexA>lexAop-rCD2::RFP, LC11-splitGAL4>UAS-CD8::GFP; nSyb-lexA>lexAop-rCD2::RFP and LC20-splitGAL4>UAS-CD8::GFP; nSyb-lexA>lexAop-rCD2::RFP three day-old females were processed for native fluorescence imaging as in (Pitman et al., 2011). In brief, brain were dissected in ice-cold 4% PFA and post-fixed in 4% PFA for 40-50 min. After 3× 20 min washes with PBST (0.01 M PBS with 0.5% TritonX) and 2× 20 min washes in PBS (0.01 M) brains were embedded in Vectashield and imaged with a 16x oil immersion lens on a Zeiss LSM 800 confocal microscope.

## References

Alexander, R.D. (1974). The evolution of social systems. Annu. Syst 5, 325–383.

Baines, R.A., Uhler, J.P., Thompson, A., Sweeney, S.T., and Bate, M. (2001). Altered electrical properties in *Drosophila* neurons developing without synaptic transmission. J. Neurosci. 21, 1523–1531.

Battesti, M., Moreno, C., Joly, D., and Mery, F. (2012). Spread of social information and dynamics of social transmission within *Drosophila* groups. Curr. Biol. 22, 309–313.

Card, G., and Dickinson, M.H. (2008). Visually mediated motor planning in the escape response of *Drosophila*. Curr. Biol. 18, 1300–1307.

Carr, C.E., and Christensen-Dalsgaard, J. (2015). Sound localization strategies in three predators. Brain. Behav. Evol. 86, 17–27.

Catania, K.C., Hare, J.F., and Campbell, K.L. (2008). Water shrews detect movement, shape, and smell to find prey underwater. Proc. Natl. Acad. Sci. 105, 571–576.

Combes, S.A., Rundle, D.E., Iwasaki, J.M., and Crall, J.D. (2012). Linking biomechanics and ecology through predator-prey interactions: flight performance of dragonflies and their prey. J. Exp. Biol. 215, 903–913.

Danchin, E., Nöbel, S., Pocheville, A., Dagaeff, A.-C., Demay, L., Alphand, M., Ranty-Roby, S., van Renssen, L., Monier, M., Gazagne, E., et al. (2018). Cultural flies: Conformist social learning in fruitflies predicts long-lasting mate-choice traditions. Science (80). 362, 1025–1030.

Faustino, A.I., Tacão-Monteiro, A., and Oliveira, R.F. (2017). Mechanisms of social buffering of fear in zebrafish. Sci. Rep. 7, 1–10.

Foster, W.A., and Treherne, J.E. (1981). Evidence for the dilution effect in the selfish herd from fish predation on a marine insect. Nature 293, 466–467.

Fotowat, H., and Gabbiani, F. (2011). Collision detection as a model for sensory-motor integration. Annu. Rev. Neurosci. 34, 1–19.

Friedel, P., Young, B.A., and Van Hemmen, J.L. (2008). Auditory localization of ground-borne vibrations in snakes. Phys. Rev. Lett. 100, 2–5.

Gibson, W.T., Gonzalez, C.R., Fernandez, C., Ramasamy, L., Tabachnik, T., Du, R.R., Felsen, P.D., Maire, M.R., Perona, P., and Anderson, D.J. (2015). Behavioral responses to a repetitive visual threat stimulus express a persistent state of defensive arousal in *Drosophila*. Curr. Biol. 25, 1401–1415.

Hamilton, W.D. (1971). Geometry for the selfish herd. J. Theor. Biol. 31, 295–311.

Handegard, N.O., Boswell, K.M., Ioannou, C.C., Leblanc, S.P., Tjøstheim, D.B., and Couzin, I.D. (2012). The dynamics of coordinated group hunting and collective information transfer among schooling prey. Curr. Biol. 22, 1213–1217.

Herberholz, J., and Marquart, G.D. (2012). Decision making and behavioral choice during predator avoidance. Front. Neurosci. 6, 1–15.

Hingee, M., and Magrath, R.D. (2009). Flights of fear: a mechanical wing whistle sounds the alarm in a flocking bird. Proc. R. Soc. B 276, 4173–4179.

Joly, J.S., Recher, G., Brombin, A., Ngo, K., and Hartenstein, V. (2016). A conserved developmental mechanism builds complex visual systems in insects and vertebrates. Curr. Biol. 26, R1001–R1009.

Kacsoh, B.Z., Bozler, J., Ramaswami, M., and Bosco, G. (2015). Social communication of predator-induced changes in *Drosophila* behavior and germ line physiology. Elife 4, 1–36.

Keleş, M.F., and Frye, M.A. (2017). Object-detecting neurons in *Drosophila*. Curr. Biol. 27, 680–687.

Lee, T., Lee, A., and Luo, L. (1999). Development of the *Drosophila* mushroom bodies: sequential generation of three distinct types of neurons from a neuroblast. Development 126, 4065–4076.

Lopes, G., Bonacchi, N., Frazão, J., Neto, J.P., Atallah, B. V., Soares, S., Moreira, L., Matias, S., Itskov, P.M., Correia, P.A., et al. (2015). Bonsai: an event-based framework for processing and controlling data streams. Front. Neuroinform. 9, 1–14.

Mortimer, B., Rees, W.L., Koelemeijer, P., and Nissen-meyer, T. (2018). Classifying elephant behaviour through seismic vibrations. Curr. Biol. 28, R547–R548.

Muijres, F.T., Elzinga, M.J., Melis, J.M., and Dickinson, M.H. (2014). Flies evade looming targets by executing rapid visually directed banked turns. Science 344, 172–177.

Murray, T.G., Zeil, J., Magrath, R.D., Murray, T.G., Zeil, J., and Magrath, R.D. (2017). Sounds of modified flight feathers reliably signal danger in a pigeon. Curr. Biol. 27, 3520–3525.

Ono, M., and Sasaki, M. (1987). Heat production by bailing in the Japanese honeybee, *Apis ceranajaponica* as a defensive behavior against the hornet, *Vespa simillima xanthoptera* (Hymenoptera: Vespidae). Experientia 43, 3–6.

Peek, M.Y., and Card, G.M. (2016). Comparative approaches to escape. Curr. Opin. Neurobiol. 41, 167–173.

Pereira, A.G., and Moita, M.A. (2016). Is there anybody out there ? Neural circuits of threat detection in vertebrates. Curr. Opin. Neurobiol. 41, 179–187.

Pereira, A.G., Cruz, A., Lima, S.Q., and Moita, M.A. (2012). Silence resulting from the cessation of movement signals danger. Curr. Biol. 22, R627–R628.

Pérez-Escudero, A., Vicente-Page, J., Hinz, R.C., Arganda, S., and de Polavieja, G.G. (2014). idTracker: tracking individuals in a group by automatic identification of unmarked animals. Nat. Methods 11.

Pitman, J.L., Huetteroth, W., Burke, C.J., Krashes, M.J., Lai, S.-L., Lee, T., and Waddell, S. (2011). A pair of inhibitory neurons are required to sustain labile memory in the *Drosophila* mushroom body. Curr. Biol. 21, 855–861.

Queiroz, H., and Magurran, A.E. (2005). Safety in numbers? Shoaling behaviour of the Amazonian red-bellied piranha. Biol. Lett. 1, 155–157.

Ramdya, P., Lichocki, P., Cruchet, S., Frisch, L., Tse, W., Floreano, D., and Benton, R. (2015). Mechanosensory interactions drive collective behaviour in *Drosophila*. Nature 519, 233–236.

Reyn, C.R. Von, Breads, P., Peek, M.Y., Zheng, G.Z., Williamson, W.R., Yee, A.L., Leonardo, A., and Card, G.M. (2014). A spike-timing mechanism for action selection. Nat. Neurosci. 17, 962–970.

Ribeiro, I.M.A., Drews, M., Bahl, A., Machacek, C., Borst, A., and Dickson, B.J. (2018). Visual projection neurons mediating directed courtship in *Drosophila*. Cell 174, 607–621.e18.

Rose, T.A., Munn, A.J., Ramp, D., and Banks, P.B. (2006). Foot-thumping as an alarm signal in macropodoid marsupials: Prevalence and hypotheses of function. Mamm. Rev. 36, 281–298.

Sanes, J.R., and Masland, R.H. (2015). The types of retinal ganglion cells: current status and implications for neuronal classification.

Sanes, J.R., and Zipursky, S.L. (2010). Design principles of insect and vertebrate visual systems. Neuron 66, 15–36.

Sarin, S., and Dukas, R. (2009). Social learning about egg-laying substrates in fruitflies. Proc. Biol. Sci. 276, 4323–4328.

Underwood, R. (1982). Vigilance behaviour in grazing african antelopes. Behaviour 79, 81–107.

Wu, M., Nern, A., Ryan Williamson, W., Morimoto, M.M., Reiser, M.B., Card, G.M., and Rubin, G.M. (2016). Visual projection neurons in the *Drosophila* lobula link feature detection to distinct behavioral programs. Elife 5, 1–43.

Zacarias, R., Namiki, S., Card, G.M., Vasconcelos, M.L., and Moita, M.A. (2018). Speed dependent descending control of freezing behavior in *Drosophila melanogaster*. Nat. Commun. 9, 3697.

Zhao, Z. dong, Chen, Z., Xiang, X., Hu, M., Xie, H., Jia, X., Cai, F., Cui, Y., Chen, Z., Qian, L., et al. (2019). Zona incerta GABAergic neurons integrate prey-related sensory signals and induce an appetitive drive to promote hunting. Nat. Neurosci. 22, 921–932.

