## Supplemental Figures 1 to 5 for "Behavioral and neuronal underpinnings of safety in numbers in fruit flies"

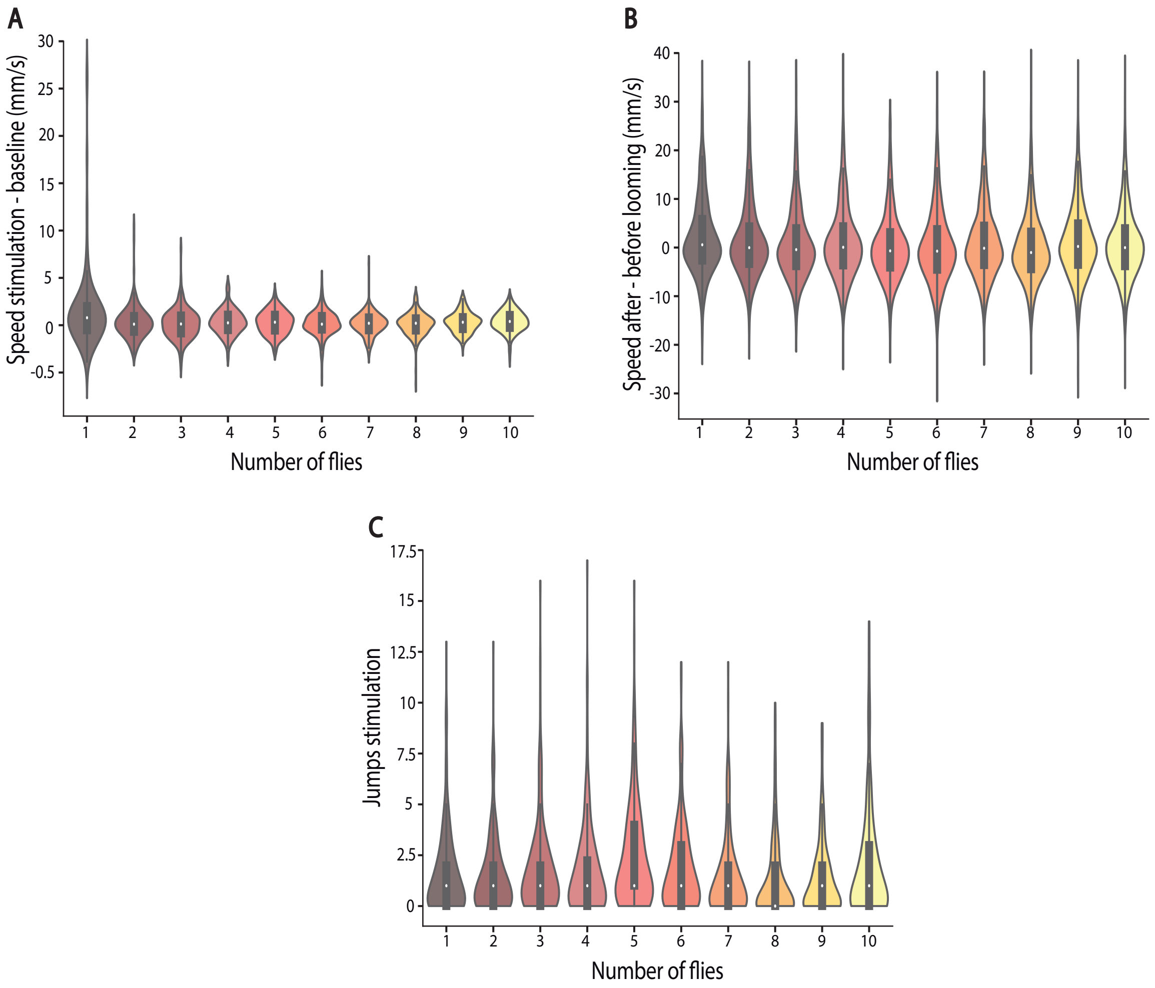


Fig. S1. Group effect on running and jumps. Related to Figure 1. A) Difference in speed between the baseline and stimulation. B) Speed differences 1 s around looming. C) Number of jumps during the stimulation.

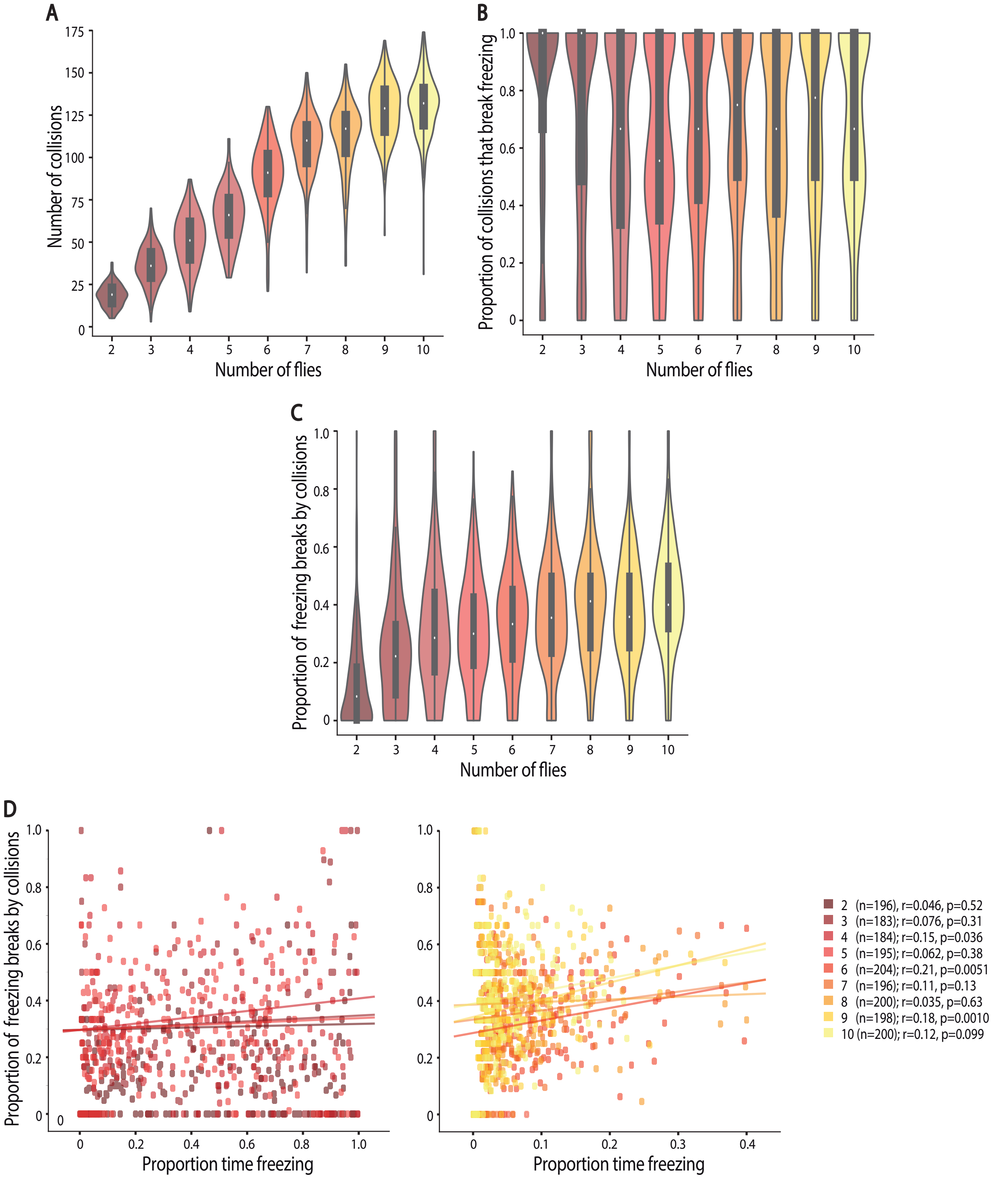


Fig. S2. The role of collisions on the group effect on freezing responses. Related to Figure 3. A-C) Violin plots (probability density of data bound to the range of possible values) with boxplots representing median values and interquartile range. A) Number of collisions throughout the experiment. Statistical comparisons between conditions presented in Table S4. B) Proportion of collisions that break freezing. Statistical comparisons between conditions presented in Table S5. C) Proportion of freezing breaks by collision. Statistical comparisons between conditions presented in Table S6. D) Correlation between the proportion of freezing breaks by collision and the proportion of time freezing for groups of 2 to 5 (left) and groups of 6 to 10 individuals (right).

The number of collisions an individual experiences increased with the increase in group size and collisions were very effective at breaking freezing. However, although the proportion of freezing breaks due to collisions increased with group size, even in groups of 10 individuals, where collisions only led to 40% IQR 31.58-53.725% of freezing breaks. Furthermore, the fact that the correlation between freezing breaks by collision and proportion of time freezing, were inexistent or very weak, further shows that collision only partly contribute to the group effect on freezing.

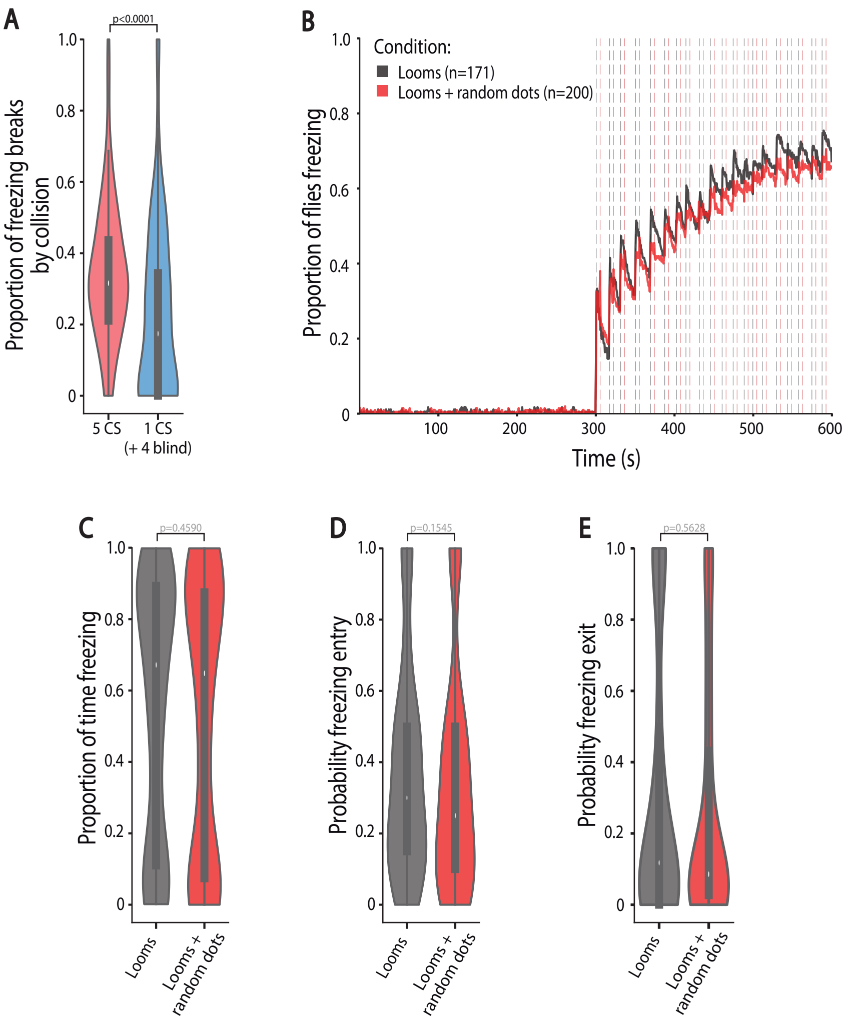


Fig. S3. Specificity of the motion cues for the group effect. Related to Figure 4. A) Effect of manipulating the motion signal with blind flies on the proportion of freezing breaks by collision. Violin plot representing the probability density of individual fly data bound to the range of possible values individual fly data, with boxplots. P-value result from Mann-Whitney test. Since the number of freezing breaks by collision decreased in groups with blind flies, the decrease in freezing observed in these groups cannot be accounted for by contacts between flies. B-E) The effect of visual disruption after looming. B) Proportion of flies freezing throughout the experiment, for individuals exposed to loomings alone or looms followed by random dots. C–E) Violin plots representing the probability density of individual fly data bound to the range of possible values individual fly data, with boxplots. C) Proportion of time spent freezing throughout the experiment. D) Probability of freezing entry in the 500 ms bin following looming presentation. E) Probability of freezing exit in the 500 ms bin before the following looming stimulus. P-values result from Kruskal-Wallis statistical analysis followed by Dunn's multiple comparisons test.

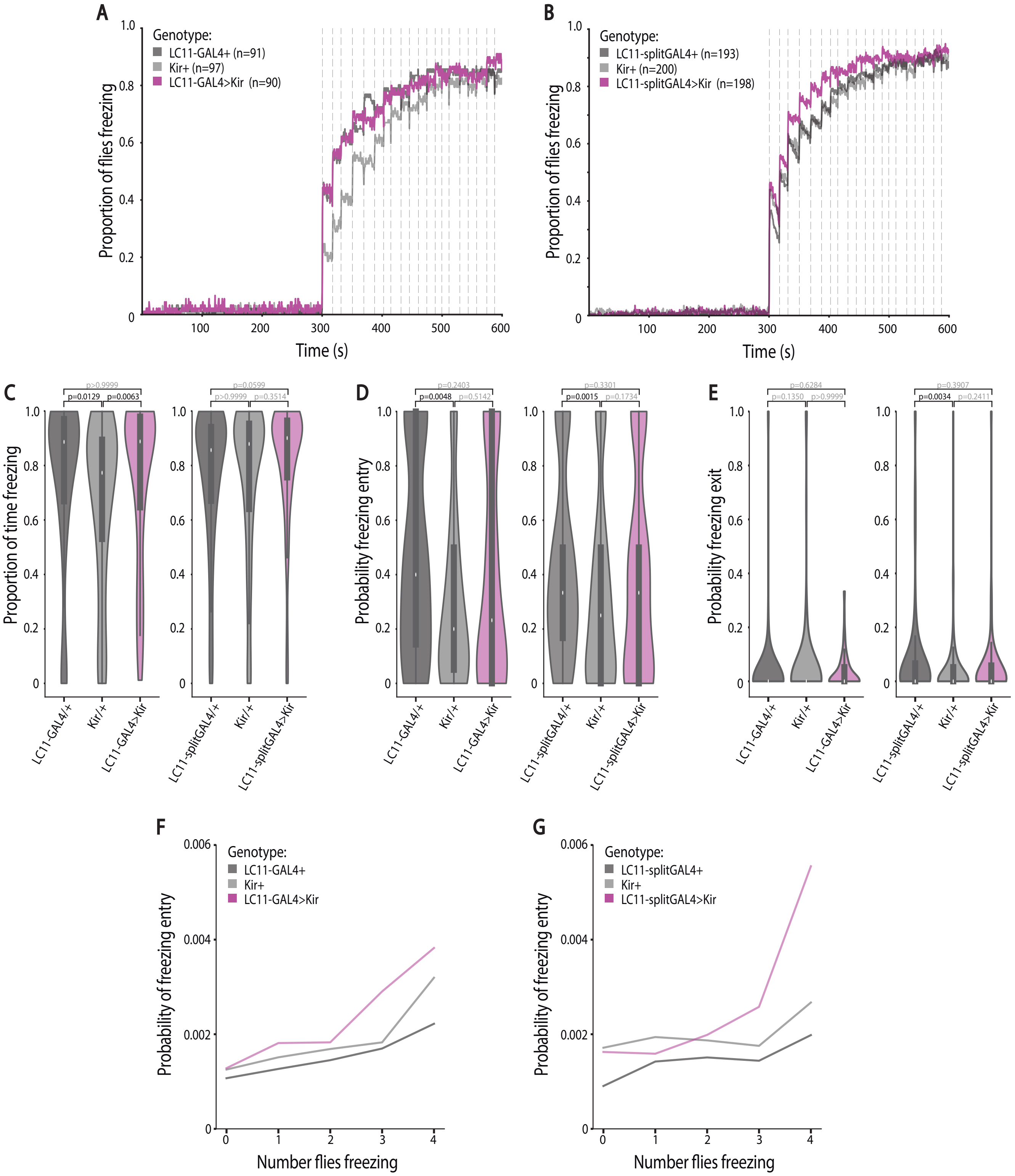


Fig. S4. Effect of manipulating lobula columnar neurons 11 (LC11) on flies tested individually and on the probability of freezing entry in groups of 5. Related to Figure 5. A–B) Proportion of flies freezing throughout the experiment, for A) *LC11-GAL4>Kir2.1* and B) *LC11-splitGAL4>Kir2.1* depicted in purple and parental controls (grey). C–E) Violin plots representing the probability density of individual fly data bound to the range of possible values, with boxplots. C) Proportion of time spent freezing throughout the experiment. D) Probability of freezing entry in the 500 ms bin following looming presentation. E) Probability of freezing exit in the 500 ms bin before the following looming stimulus. P-values result from Kruskal-Wallis statistical analysis followed by Dunn's multiple comparisons test. F-G) Probability of freezing entry at time t as a function of the number of other flies freezing at t-1 (see methods) in groups of 5 for F) *LC11-GAL4>Kir2.1* and G) *LC11-splitGAL4>Kir2.1* depicted in purple and parental controls (grey). The observation that LC11-silenced flies show a bigger effect of surrounding flies freezing is likely due to the enhanced freezing responses in this group of flies.

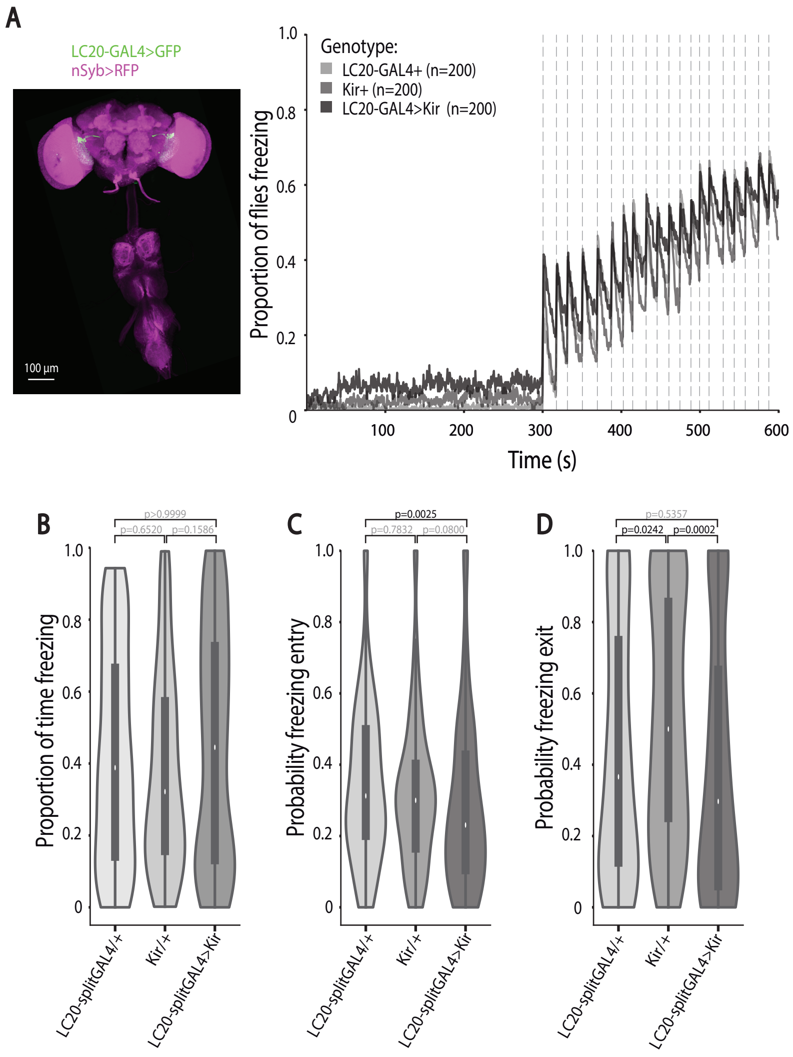


Fig. S5. Manipulating lobula columnar neurons 20 (LC20). Related to Figure 5. A) Anatomy (scale bar, 100 µm) and proportion of flies freezing throughout the experiment in groups of 5, for *LC20-splitGAL4>Kir2.1* in dark grey, compared to parental controls (light grey). B–D) Violin plots representing the probability density of individual fly data bound to the range of possible values, with boxplots. B) Proportion of time spent freezing throughout the experiment. C) Probability of freezing entry in the 500 ms bin following looming presentation. D) Probability of freezing exit in the 500 ms bin before the following looming stimulus. P-values result from Kruskal-Wallis statistical analysis followed by Dunn's multiple comparisons test.
