## Supplemental Tables 1 to 6 for "Behavioral and neuronal underpinnings of safety in numbers in fruit flies"

Supplemental Table 1. Statistical comparisons of the time each fly spent freezing when tested individually and in groups of up to 10 individuals (p-values for Kruskall-Wallis followed by Dunn's multiple comparisons test).

| | 2 | 3 | 4 | 5 | 6 | 7 | 8 | 9 | 10 |
|---|---|---|---|---|---|---|---|---|---|
| 1 | 0.0272 | 0.0229 | 0.0009 | 0.0001 | <0.0001 | <0.0001 | <0.0001 | <0.0001 | <0.0001 |
| 2 | | >0.9999 | >0.9999 | >0.9999 | <0.0001 | <0.0001 | <0.0001 | <0.0001 | <0.0001 |
| 3 | | | >0.9999 | >0.9999 | <0.0001 | <0.0001 | <0.0001 | <0.0001 | <0.0001 |
| 4 | | | | >0.9999 | <0.0001 | <0.0001 | <0.0001 | <0.0001 | <0.0001 |
| 5 | | | | | <0.0001 | <0.0001 | <0.0001 | <0.0001 | <0.0001 |
| 6 | | | | | | 0.1189 | 0.5697 | <0.0001 | 0.0004 |
| 7 | | | | | | | >0.9999 | >0.9999 | >0.9999 |
| 8 | | | | | | | | 0.5132 | >0.9999 |
| 9 | | | | | | | | | >0.9999 |

Supplemental Table 2. Statistical comparisons of the probability each fly started freezing after a looming stimulus when tested individually and in groups of up to 10 individuals (p-values for Kruskall-Wallis followed by Dunn's multiple comparisons test).

| | 2 | 3 | 4 | 5 | 6 | 7 | 8 | 9 | 10 |
|---|---|---|---|---|---|---|---|---|---|
| 1 | >0.9999 | >0.9999 | >0.9999 | >0.9999 | >0.9999 | <0.0001 | 0.0007 | <0.0001 | <0.0001 |
| 2 | | >0.9999 | >0.9999 | 0.5177 | >0.9999 | <0.0001 | 0.0114 | <0.0001 | <0.0001 |
| 3 | | | >0.9999 | >0.9999 | 0.1128 | <0.0001 | <0.0001 | <0.0001 | <0.0001 |
| 4 | | | | >0.9999 | >0.9999 | <0.0001 | 0.0011 | <0.0001 | <0.0001 |
| 5 | | | | | 0.0029 | <0.0001 | <0.0001 | <0.0001 | <0.0001 |
| 6 | | | | | | 0.0113 | >0.9999 | 0.0003 | 0.0009 |
| 7 | | | | | | | >0.9999 | >0.9999 | >0.9999 |
| 8 | | | | | | | | >0.9999 | >0.9999 |
| 9 | | | | | | | | | >0.9999 |

Supplemental Table 3. Statistical comparisons of probability each fly stopped freezing before the following looming stimulus when tested individually and in groups of up to 10 individuals (p-values for Kruskall-Wallis followed by Dunn's multiple comparisons test).

| | 2 | 3 | 4 | 5 | 6 | 7 | 8 | 9 | 10 |
|---|---|---|---|---|---|---|---|---|---|
| 1 | <0.0001 | <0.0001 | <0.0001 | <0.0001 | <0.0001 | <0.0001 | <0.0001 | <0.0001 | <0.0001 |
| 2 | | >0.9999 | >0.9999 | 0.0395 | <0.0001 | <0.0001 | <0.0001 | <0.0001 | <0.0001 |
| 3 | | | >0.9999 | 0.3299 | <0.0001 | <0.0001 | <0.0001 | <0.0001 | <0.0001 |
| 4 | | | | >0.9999 | <0.0001 | <0.0001 | <0.0001 | <0.0001 | <0.0001 |
| 5 | | | | | <0.0001 | <0.0001 | <0.0001 | <0.0001 | <0.0001 |
| 6 | | | | | | >0.9999 | >0.9999 | >0.9999 | >0.9999 |
| 7 | | | | | | | >0.9999 | >0.9999 | >0.9999 |
| 8 | | | | | | | | >0.9999 | >0.9999 |
| 9 | | | | | | | | | >0.9999 |

Supplemental Table 4. Statistical comparisons of the number of collisions each fly experienced throughout the experiment when tested in groups of 2 to 10 individuals (p-values for Kruskall-Wallis followed by Dunn's multiple comparisons test).

| | 3 | 4 | 5 | 6 | 7 | 8 | 9 | 10 |
|---|---|---|---|---|---|---|---|---|
| 2 | 0.0014 | <0.0001 | <0.0001 | <0.0001 | <0.0001 | <0.0001 | <0.0001 | <0.0001 |
| 3 | | 0.0664 | <0.0001 | <0.0001 | <0.0001 | <0.0001 | <0.0001 | <0.0001 |
| 4 | | | 0.1555 | <0.0001 | <0.0001 | <0.0001 | <0.0001 | <0.0001 |
| 5 | | | | <0.0001 | <0.0001 | <0.0001 | <0.0001 | <0.0001 |
| 6 | | | | | 0.0001 | <0.0001 | <0.0001 | <0.0001 |
| 7 | | | | | | >0.9999 | <0.0001 | <0.0001 |
| 8 | | | | | | | 0.0013 | <0.0001 |
| 9 | | | | | | | | >0.9999 |

Supplemental Table 5. Statistical comparisons of the proportion of collisions that led to freezing exit per fly when tested groups of 2 to 10 individuals (p-values for Kruskall-Wallis followed by Dunn's multiple comparisons test).

| | 3 | 4 | 5 | 6 | 7 | 8 | 9 | 10 |
|---|---|---|---|---|---|---|---|---|
| 2 | 0.1090 | <0.0001 | <0.0001 | <0.0001 | 0.0013 | <0.0001 | 0.0120 | 0.0002 |
| 3 | | 0.3963 | 0.0126 | >0.9999 | >0.9999 | 0.2947 | >0.9999 | >0.9999 |
| 4 | | | >0.9999 | >0.9999 | >0.9999 | >0.9999 | >0.9999 | >0.9999 |
| 5 | | | | >0.9999 | 0.4059 | >0.9999 | 0.0732 | >0.9999 |
| 6 | | | | | >0.9999 | >0.9999 | >0.9999 | >0.9999 |
| 7 | | | | | | >0.9999 | >0.9999 | >0.9999 |
| 8 | | | | | | | 0.9607 | >0.9999 |
| 9 | | | | | | | | >0.9999 |

Supplemental Table 6. Statistical comparisons of the proportion of freezing breaks by collision per fly when tested in groups of 2 to 10 individuals (p-values for Kruskall-Wallis followed by Dunn's multiple comparisons test).

| | 3 | 4 | 5 | 6 | 7 | 8 | 9 | 10 |
|---|---|---|---|---|---|---|---|---|
| 2 | <0.0001 | <0.0001 | <0.0001 | <0.0001 | <0.0001 | <0.0001 | <0.0001 | <0.0001 |
| 3 | | 0.0636 | 0.0258 | <0.0001 | <0.0001 | <0.0001 | <0.0001 | <0.0001 |
| 4 | | | >0.9999 | >0.9999 | 0.3180 | 0.0002 | 0.0275 | <0.0001 |
| 5 | | | | >0.9999 | 0.5244 | 0.0003 | 0.0494 | <0.0001 |
| 6 | | | | | >0.9999 | 0.1155 | >0.9999 | 0.0065 |
| 7 | | | | | | >0.9999 | >0.9999 | 0.1938 |
| 8 | | | | | | | >0.9999 | >0.9999 |
| 9 | | | | | | | | >0.9999 |
